# Combining population genomics and forward simulations to investigate stocking impacts: A case study of Muskellunge (*Esox masquinongy*) from the St. Lawrence River basin

**DOI:** 10.1101/363283

**Authors:** Quentin Rougemont, Anne Carrier, Jeremy Leluyer, Anne-Laure Ferchaud, John M. Farrell, Daniel Hatin, Philippe Brodeur, Louis Bernatchez

**Affiliations:** Département de biologie, Institut de Biologie Intégrative et des Systèmes (IBIS), Université Laval, G1V 0A6, Québec, Canada; IFREMER, Unité Ressources Marines en Polynésie, Centre Océanologique du Pacifique - Vairao - BP 49 - 98179 Taravao - Tahiti - Polynésie Française; Department of Environmental and Forest Biology, State University of New York, College of Environmental Science and Forestry, 13210, Syracuse, New York, USA; Ministère des Forêts, de la Faune et des Parcs, Direction de la Gestion de la Faune Estrie-Montréal-Montérégie-Laval, 201 Place Charles-Le Moyne, Longueuil, Québec J4K 2T5, Canada; Ministère des Forêts, de la Faune et des Parcs, Direction de la gestion de la faune de la Mauricie et du Centre-du-Québec, 100 rue Laviolette, bureau 207, Trois-Rivières, G9A 5S9, Canada

**Keywords:** admixture, gene flow, forward simulations, stocking, genomic, *Esox masquinongy*

## Abstract

Understanding the genetic and evolutionary impacts of fish stocking on wild populations has long been of interest as negative consequences such as reduced fitness and loss of genetic resources are commonly reported. Nearly five decades of extensive stocking of over a million Muskellunge (*Esox masquinongy*) in the Lower St. Lawrence River (Québec, Canada) was implemented by managers in an attempt to sustain a fishery. We investigated the effect of stocking on this native species’ genetic structure and allelic diversity in the St. Lawrence river and its tributaries as well as several stocked inland lakes. Using Genotype-By-Sequencing (GBS), we genotyped 643 individuals representing 22 sampling sites and combined this information with forward simulations to investigate the genetic consequences of stocking. Individuals native to the St. Lawrence watershed were genetically divergent from the sources used for stocking and both the St. Lawrence tributaries and inland lakes were also naturally divergent from the main stem. Empirical data and simulations revealed weak effects of stocking on admixture patterns within the St. Lawrence despite intense stocking in the past, whereas footprints of admixture were higher in the smaller stocked populations from tributaries and lakes. Altogether, our data suggests that selection against introgression has been relatively efficient within the large St. Lawrence River. In contrast, the smaller populations from adjacent tributaries and lakes still displayed stocking related admixture which apparently resulted in higher genetic diversity, suggesting that, while stocking stopped at the same time, its impact remained higher in these populations. Finally, the origin of populations from inland lakes that were established by stocking confirmed their close affinity with these source populations. This study illustrates the benefit of combining extensive genomic data with forward simulations for improved inferences regarding outcomes of population enhancement by stocking, as well as its relevance for fishery management decision making.

## Introduction

Many species and populations are undergoing steep declines in abundance due to a globally changing environment and to overexploitation (Allendorf, 2017). As a result, supplementation programs continue to be extensively applied to enhance natural populations, both for conservation purposes and sustaining exploitation, for instance in fisheries and forestry sectors (Laikre, Schwartz, Waples, Ryman, & GeM Working Group, 2010). This is particularly true for fish that are subjected to commercial and recreational exploitation (Dunham, 2011) and also for species sometimes transferred over large distances to supplement genetically and ecologically divergent populations. As a consequence, understanding how supplementation practices from non-indigenous sources might impact the ecological and genetic integrity as well as the evolutionary potential of wild populations remains a major concern (Araki, Cooper, & Blouin, 2007, 2009; Laikre & Ryman, 1996; Ryman & Laikre, 1991; Waples, Hindar, Karlsson, & Hard, 2016). From a genetic standpoint, potential impacts of stocking include: reduction in genetic diversity and effective population size due to the Ryman-Laikre effect (i.e. the use of a low number of individual for reproduction in supportive breeding programs Laikre & Ryman, 1996; Ryman & Laikre, 1991), genetic homogenization of wild populations (Araki & Schmid, 2010; Eldridge, Myers, & Naish, 2009; Eldridge & Naish, 2007; Lamaze, Sauvage, Marie, Garant, & Bernatchez, 2012; Perrier, Guyomard, Bagliniere, Nikolic, & Evanno, 2013) and ultimately, the loss of local adaption (Araki et al., 2009; Ford, 2002; Lynch & O’Hely, 2001). Loss of local adaptation may stem from the fact that selected traits in captive environments or any foreign stocking source may differ from those providing the highest fitness in the local environment (Fraser et al. 2018). Following admixture (impacting genome-wide structure), and eventual introgression (in which allelic variants can be transferred from one differentiated population to another), such differences in selective values, together with higher genetic load in stocked populations can lead to outbreeding depression resulting from disruption of co-adapted gene complexes or from the breakup of epistatic interactions (Lynch, 1991; Tallmon, Luikart, & Waples, 2004). Such results have been documented in numerous occasions (e.g. Allendorf, Leary, Spruell, & Wenburg, 2001; Edmands, 2007; Le Cam, Perrier, Besnard, Bernatchez, & Evanno, 2015), which raise concerns related to the consequences of stocking on the long term maintenance of wild populations (Létourneau et al., 2018).

While the negative effects of admixture and introgression have been extensively documented in various salmonids species (Finnegan & Stevens, 2008; Hansen, Fraser, Meier, & Mensberg, 2009; Létourneau et al., 2018; Perrier, Baglinière, & Evanno, 2013; Perrier, Guyomard, et al., 2013), this has been much less investigated in other taxonomic groups. Here we address this general question for the Muskellunge *Esox masquinongy*, an Esocidae which is the sister family of all salmonids (Rondeau et al., 2014). Muskellunge is distributed throughout temperate rivers and lakes of Northeastern America (Crossman, 1978), where it often occurs in sympatry withthe Northern Pike *Esox lucius* a closely related species. It reaches sexual maturity between 5 to 7 years for males and 6 to 8 years for females (Farrell et al., 2007) and can grow up to more than 1.5 m and 20 kg, which makes it a highly prized species for recreational trophy fishing. Given its typically low population density (~ < 1.0 fish/ha; Cloutier, 1987; Simonson, 2010), relatively little is known regarding the species biology, genetic diversity and population dynamics (Crane et al., 2015; Kapuscinski, Sloss, & Farrell, 2013).

In many places, Muskellunge have undergone pronounced population decline due to various factors including over-harvesting, habitat degradation and pollution (Mongeau et Massé, 1976; Crossman, 1978, Farrell et al., 2007; Whillans, 1979; Brodeur et al. 2013), and more recently as a consequence of viral hemorrhagic septicemia and round goby introduction (Farrell et al. 2017). As a result, numerous water bodies have been subjected to extensive stocking to rehabilitate native populations, but also to introduce the species in habitats it never occupied for recreational purposes and to control invasive fish species (Wingate, 1986; Crane et al., 2015; Jennings et al., 2010; Kapuscinski, Belonger, Fajfer, & Lychwick, 2007). The genetic structure that prevailed before stocking and its effects on the genetic integrity of wild populations have been rarely investigated in Muskellunge. Most studies thus far pertaining to the species genetic structure used the few available microsatellite markers and primarily focused on the potential effect of stocking and genetic divergence in the Laurentian Great Lakes and Upper St. Lawrence River (Kapuscinski et al., 2013; Miller, Mero, & Younk, 2009, 2012; Turnquist et al., 2017; Wilson, Liskauskas, & Wozney, 2016). These studies were useful in revealing a pronounced genetic differentiation among weakly connected populations at small spatial scales, as well as a generally reduced genetic diversity putatively associated to recent bottlenecks and/or strong genetic drift in populations of small effective size and restricted dispersal due to a tendency toward spawning site fidelity (Crossman, 1990; Jennings, Hatzenbeler, & Kampa, 2011; Margenau, 1994; Miller, Kallemeyn, & Senanan, 2001). Moreover, the impact of stocking has been documented in several populations where modest levels of admixture and a partial homogenization of population genetic structure have been reported (Scribner et al., 2015). In contrast, no study has documented the species population structure further east, in the Lower St. Lawrence River basin of Québec (Canada), downstream of the Great Lakes where Muskellunge populations are native. Moreover, no population genomics studies using thousands of markers distributed throughout the genome has been performed on this species to date. As in the rest of its distribution range, the species in the St. Lawrence area underwent a steep decline attributed to habitat loss (Robittaille & Cotton, 1992) and commercial overfishing that occurred until 1936 (Dymond 1939; Crossman 1986). Therefore, the lower St. Lawrence had been supplemented by over one million of young Muskellunge from 1951 to 1998 (Vézina, 1977; Vincent et Legendre 1974; Mongeau et al. 1980; Dumont, 1991; Brodeur et al. 2013; De La Fontaine unpublished). In particular, during the time period spanning 1951-1965, fish from both New York State (Chautauqua Lake, Ohio River basin) and Ontario’s Kawartha Lakes (represented in our study by Pigeon Lake) were transferred to Lachine hatcheries in Québec to support a massive stocking program in the St. Lawrence River (Dumont, 1991; De La Fontaine, unpublished). Muskellunge from these two lakes not only come from a very distinct geographic area, but also are genetically different from each other and from those of the Great Lakes (Koppelman & Phillip, 1986; Turnquist et al., 2017). Fish from those foreign sources were also stocked in numerous lakes including Lake Joseph and Lake Tremblant which themselves became widely used as a brood source from 1965 for hatcheries until the end of stocking in 1998 (Vézina, 1977; Vincent et Legendre, 1974; Dumont 1991; De La Fontaine, unpublished). Besides massive stocking in those waterbodies, Muskellunge were also introduced in several lakes and rivers where it never occurred before in Québec. Details on the history of stocking and translocations are provided in supplementary Fig S1.

Besides traditional population genetics, an improved understanding of the possible effects of admixture following stocking can be achieved by the application of individually-based forward simulations (Hoban, 2014; Hoban, Bertorelle, & Gaggiotti, 2012). Simulations may be used to compare the simulated outcomes of population divergence following stocking resulting from a variety of demographic scenarios (e.g. in terms of population size and differential mortality) in order to identify the one that best explains empirically documented patterns (Guillaume & Rougemont, 2006; Hernandez, 2008; Lawrie, 2017; Messer, 2013). Yet, such methods have rarely been applied to investigate the effect of stocking on the genetic composition of wild fish populations (but see Perrier et al., 2013). It is also possible to generate increasingly complex and biologically realistic simulations that can be compared to an empirical dataset. Such simulations can aid in understanding the efficacy of supplementation and how it may generate admixture, while also helping to understand the limits of using simple clustering tools (e.g. Alexander, Novembre, & Lange, 2009; Pritchard, Stephens, & Donnelly, 2000) to accurately detect the genome wide effect of admixture on local ancestry.

In this study, we combined empirical population genomics using a Genotype-By-Sequencing (GBS) approach to forward simulations in order to document the extent and patterns of genetic admixture that resulted from past stocking events in the St. Lawrence River watershed. A total 643 fish were collected at 22 locations in the St. Lawrence watershed including its major tributaries and adjacent inland lakes. The four main historical stocking sources used in Québec were also sampled. In addition, a single lake for which there is no record of stocking and therefore presumed to be the only known fully native lacustrine Muskellunge population in Québec was included in the analysis (Table 1, Fig 1). Following the more conventional analyses of population structure based on clustering and measures of genetic differentiation, we matched our empirical data to simplified scenarios of divergence and admixture generated using forward simulations. With this dataset we aimed at *i*) exploring the effect of nearly five decades of stocking on wild populations and documenting the fine scale population genetic diversity and structure of the Muskellunge in the St. Lawrence River watershed, *ii*) comparing the expected patterns of admixture from simplified demographic simulations with empirical observations, and *iii*) using the information to define management units and make recommendations pertaining to stocking practices.

**Table 1:** Genetic diversity parameters. Code = acronyms for each sampling site, n = number of individuals genotyped, Ho = observed heterozygosity, Hs = gene diversity, P = Proportion of polymorphic loci, Ne = effective population size with 95% CI obtained from jackknifing over individuals. Effective population size was computed by merging the major group within the St. Lawrence together. Similarly, all populations from the same stocking group (JOS, TRE and FRO) were merged together. Such merging was performed in order to increase the sample size. Effective population size could not be estimated for Traverse Lake when implementing a correction for linkage and resulted in an estimated of Ne = 3 [95%IC = 2-4] without correction.

| Sampling location | code | type | n | Ho | Hs | P | Pi | Ne [95% CI] |
| --- | --- | --- | --- | --- | --- | --- | --- | --- |
| **Upper Saint Lawrence** |  |  |  |  |  |  |  |  |
| Thousand Island National Park | TIN | stocked | 18 | 0.154 | 0.160 | 0.181 | 0.0033 | 362 [150-inf ] |
| St. Lawrence Lake | LSW | stocked | 15 | 0.169 | 0.174 | 0.177 | 0.0034 | 669 [493-823] |
| **Lower Saint Lawrence and inland lake** |  |  |  |  |  |  |  |  |
| Saint-Louis Lake | LSL | stocked | 55 | 0.099 | 0.101 | 0.305 | 0.0030 | 669 [493-823] |
| Saint-François Lake | LSF | stocked | 61 | 0.102 | 0.104 | 0.286 | 0.0030 | 669 [493-823] |
| Saint-Pierre Lake | LSP | stocked | 57 | 0.101 | 0.102 | 0.298 | 0.0030 | 669 [493-823] |
| Contrecoeur (St. Lawrence River) | MSC | stocked | 16 | 0.163 | 0.168 | 0.176 | 0.0034 | 669 [493-823] |
| Boucherville (St. Lawrence River) | MSG | stocked | 19 | 0.144 | 0.150 | 0.209 | 0.0033 | 669 [493-823] |
| Varennes (St. Lawrence River) | MSV | stocked | 16 | 0.161 | 0.165 | 0.189 | 0.0034 | 669 [493-823] |
| **Tributaries** |  |  |  |  |  |  |  |  |
| des Deux-Montagnes Lake | LDM | stocked | 56 | 0.110 | 0.114 | 0.287 | 0.0030 | 146 [80-298] |
| De l'Achigan River | ACH | stocked | 25 | 0.149 | 0.154 | 0.170 | 0.0029 | 17 [11-178] |
| Chaudière River | CHD | Introduced | 15 | 0.196 | 0.198 | 0.150 | 0.0032 | 375 [84-inf] |
| Saint-Maurice River | MAU | stocked | 26 | 0.172 | 0.174 | 0.161 | 0.0031 | 45 [25-77] |
| Maskinongé Lake | MSK | stocked | 25 | 0.131 | 0.136 | 0.236 | 0.0032 | 52 [24 - 145] |
| Ottawa River | OTT | stocked | 7 | 0.255 | 0.277 | 0.110 | 0.0035 | 45 [17- inf] |
| Yamaska River | YAM | stocked | 16 | 0.180 | 0.183 | 0.137 | 0.0030 | 62 [29 - 129] |
| **Stocking Sources** |  |  |  |  |  |  |  |  |
| Chautauqua Lake | CHQ | source | 41 | 0.129 | 0.129 | 0.210 | 0.0028 | 137 [95 – 175] |
| Pigeon Lake | PIG | source | 27 | **0.088** | 0.086 | 0.152 | 0.0019 | 79 [36-178] |
| Joseph Lake | JOS | source | 30 | 0.139 | 0.142 | 0.190 | 0.0029 | 137 [95 – 175] |
| Tremblant Lake | TRE | source | 29 | 0.152 | 0.157 | 0.160 | 0.0029 | 137 [95 – 175] |
| **Stocked lake** |  |  |  |  |  |  |  |  |
| Champlain Lake | CHP | stocked | 23 | 0.165 | 0.174 | 0.159 | 0.0031 | 14 [6-31] |
| Frontière Lake | FRO | Introduced | 31 | 0.145 | 0.148 | 0.170 | 0.0029 | 137 [95 – 175] |
| **Unstocked lake** |  |  |  |  |  |  |  |  |
| Traverse Lake | TRA | wild | 35 | 0.070 | 0.066 | 0.154 | 0.0016 | NA [NA] |

**Figure 1.**
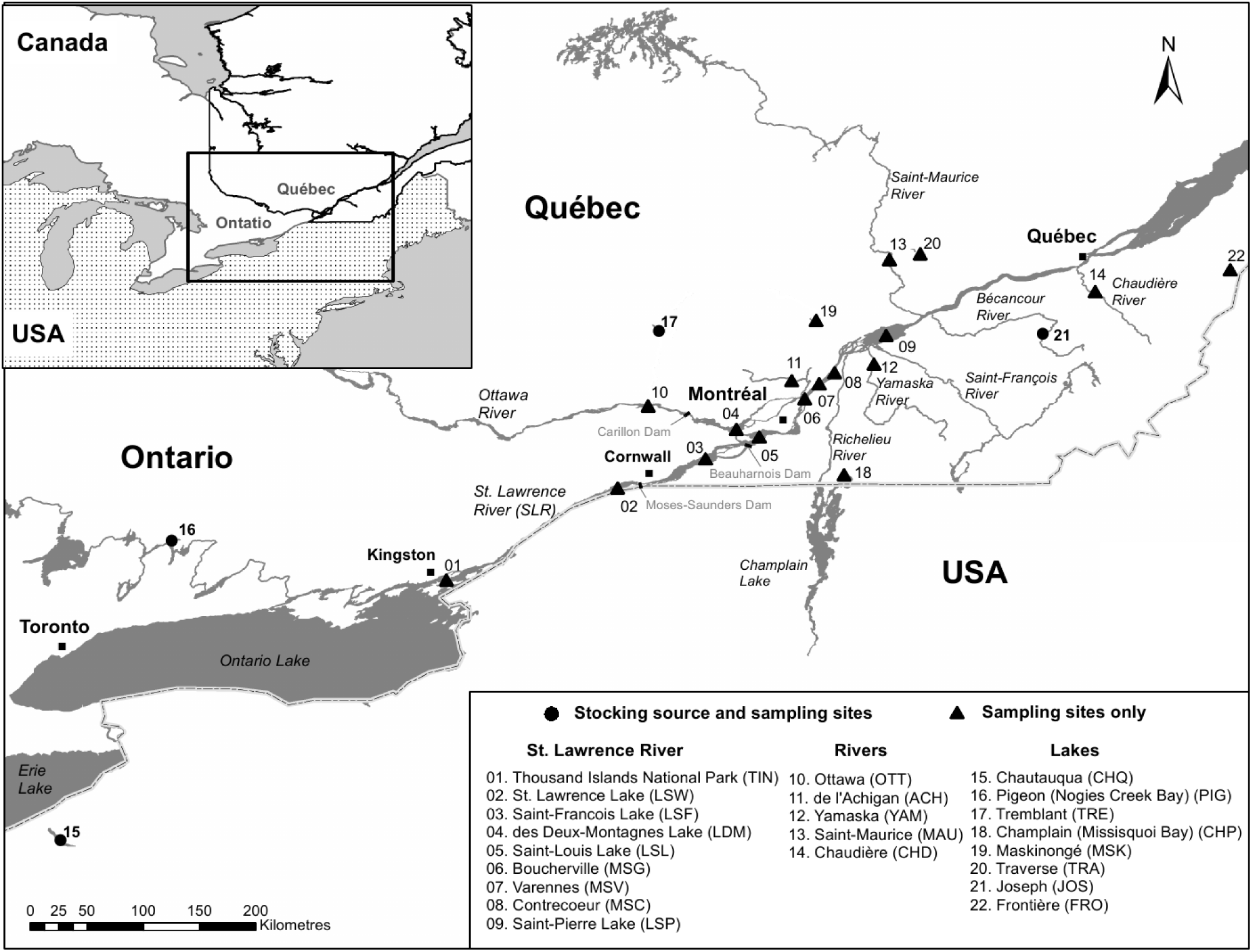
Map showing sampling locations. Two major physical barriers are present on the St. Lawrence system: Moses-Saunders and Beauharnois-Les Cèdres hydropower dams located at the upstream and downstream end of Lake Saint-Françis, respectively.

## Methods

### Sampling

A total of 662 Muskellunge were sampled from 22 sites within the St. Lawrence River drainage, mainly from Québec (Canada) with a median sample size of 24 individuals per site (Fig 1, table 1). This sampling represents the Upper and Lower St. Lawrence River, its main tributaries as well as inland lakes populations. Two sampling sites (Chautauqua Lake, CHQ, and Pigeon lake, PIG) represent the original stocking source used from 1951 to 1965 and two more recent secondary sources (Lake Joseph, JOS, 1965-1986, and Lake Tremblant, TRE, 1986-1997; both initially stocked with the CHQ and PIG source) were included in the baseline. Of all sampling locations, to the best of our knowledge, only Traverse Lake (TRA) has no record of past stocking and is presumed to be native. Most samples (fin clips preserved in 95% ethanol) were mainly obtained from catch & release captures of professional fishing guides and anglers between 2010 and 2015. Samples were also collected by the Ministère des Forêts, de la Faune et des Parcs du Québec (MFFP), the Ontario Ministry of Natural Resources and Forestry (MNRF), and the New York Department of Environmental Conservation. A graphical representation of the major stocking operations performed in Québec is presented in Supplementary Fig S1.

### 2.2 Molecular methods and SNP calling

A salt-extraction protocol adapted from (Aljanabi & Martinez, 1997) was used to extract genomic DNA. Sample quality and concentration were checked on 1% agarose gels and a NanoDrop 2000 spectrophotometer (Thermo Scientific). Concentration of DNA was normalized to 20 ng/μl. Libraries were constructed following a double-digest RAD (restriction site-associated DNA sequencing; Andrews, Good, Miller, Luikart, & Hohenlohe, 2016) protocol modified from Mascher, Wu, Amand, Stein, and Poland (2013). Genomic DNA was digested with two restriction enzymes (*Pst*I and *Msp*I) by incubating at 37°C for two hours followed by enzyme inactivation by incubation at 65°C for 20 min. Sequencing adaptors and a unique individual barcode were ligated to each sample using a ligation master mix including T4 ligase. The ligation reaction was completed at 22°C for 2 hours followed by 65°C for 20 min to inactivate the enzymes. Samples were pooled in multiplexes of 48 individuals, insuring that individuals from each sampling location were sequenced as part of at least six different multiplexes to avoid pool effects. Libraries were size-selected using a BluePippin prep (Sage Science), amplified by PCR and sequenced on the Ion Proton P1v2 chip. Ninety-six individuals were sequenced per chip. Each library was sequenced on one Ion Proton chip which generated approximately 80 million reads. Based on the number of reads for each individually barcoded sample on this first chip, the prepared DNA was re-pooled into a new library where the representation of samples with low reads counts was increased. This new library was then sequenced on two more Ion Proton chips, thus normalizing the number of reads per sample.

### Bioinformatics

Barcodes were removed using cutadapt (Martin, 2011) and trimmed to 80 pb and allowing for an error rate of 0.2. They were then demultiplexed using the ‘process_radtags’ module of Stacks v1.44 (Catchen, Hohenlohe, Bassham, Amores, & Cresko, 2013) and aligned to the Northern Pike (*Esox lucius*) reference genome Eluc_V3 (Rondeau et al., 2014) using bwa-mem (Li et al., 2009) with default parameters. Here, 19 individuals randomly distributed across sampling sites with less than 2.5 million reads were removed so that 643 individuals out of the 662 were kept for all subsequent analysis. Then, aligned reads were processed with Stacks v.1.44 for SNPs calling and genotyping. The ‘pstacks’ module was used with a minimum depth of 3 and up to 3 mismatches were allowed in the catalog creation. We then ran the ‘populations’ module to produce a vcf file that was further filtered using a custom python script. SNPs were kept if they displayed a read depth higher than 5, and were present in a least 70 % in each sampling location and did not show heterozygosity higher than 0.60 to control for paralogs and HWE disequilibrium. The resulting vcf file comprised 16,266 SNPs spread over 11,458 loci and represented the least stringent dataset used to estimate basic diversity parameters. This vcf file was then subsampled to meet the different assumptions (in particular the use of unlinked SNPs) of the models underlying the different population genetic software used below. We avoided using minor allele frequency MAF threshold to obtain a true site frequency spectrum and unbiased estimates of diversity indices. Indeed, MAF threshold generates biases that are not well accounted for in many of the analyses performed below (Guillot & Foll, 2009). A haplotype file was also exported using the ‘populations’ module in Stacks and used to estimate nucleotide diversity (Tajima, 1989) and to estimate structure in FineRADstructure analysis.

### Genetic diversity analysis

We first used Hierfstat (Goudet, 2005) to compute patterns of observed heterozygosity and gene diversity after excluding monomorphic markers from each sampling location. Second, we computed the proportion of polymorphic loci in each sampling location in R. We estimated contemporary effective population size (Ne) based on linkage disequilibrium principles and using the LDNe method (Waples & Do, 2008) implemented in NeEstimator (Do et al., 2014). We kept one single SNP by loci, and filtered the dataset for physical LD using plink (Chang et al. 2015). We used windows of 50 SNPs shifted by 5 SNPs each iteration and removed any SNP with a variation inflation factor greater than 2. We then removed non polymorphic loci in each population as well as singletons. Effective population size was computed after merging sampling locations into their major genetic clusters identified by the different clustering analyses described below and by applying equations from table 2 of Waples (2006) after removing all Burrows estimates for pairs of SNPs located on the same chromosome. Finally, we estimated nucleotide diversity (Tajima, 1989) at the haplotype level using mscalc (Ross-Ibarra, Tenaillon, & Gaut, 2009, Ross-Ibarra et al., 2008, Roux et al., 2011).

**Table 2.**
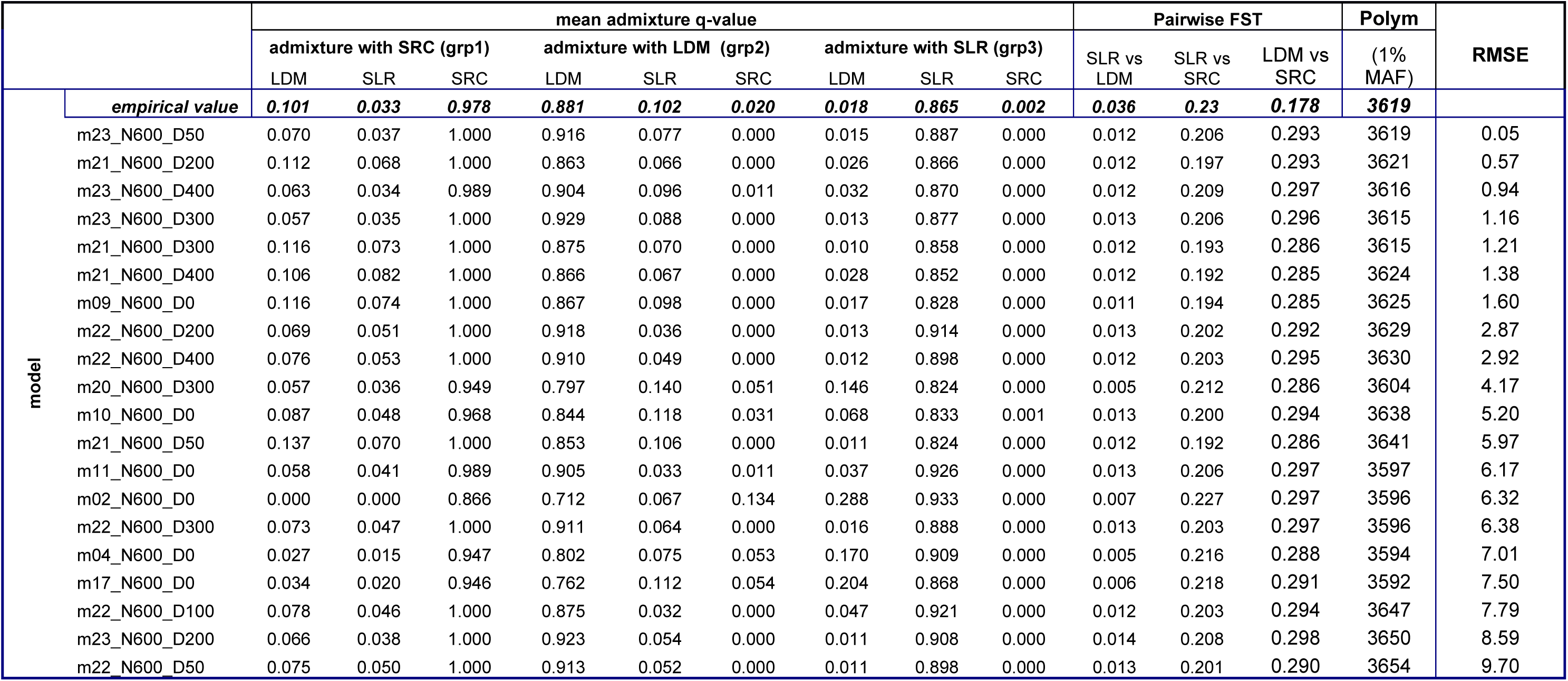
Simulation results for the 20 models showing the lowest RMSE. Abbreviation: SRC = Source population used for stocking. Given the shared patterns of ancestry between the populations used for stocking (i.e. Chautauqua, Joseph and Tremblant lakes), individuals were merged together and modeled as a single unit. Given that no sign of ancestry from Pigeon Lake was found in the St. Lawrence (SLR) or Deux-Montagnes L. (LDM), this source was not included here. Based on our empirical results, the migration rate (m) was set 1.5 times higher in LDM than in SLR in which we included LSL, LSF, LSP, MSC, MSG, and MSV. Admixture and *F_ST_* values in bold are those observed in our empirical data. The group 1 (grp1), group 2 (grp2) and group 3 (grp3) represent admixture membership probability of individuals from LDM, SLR and SRC respectively for K = 3. Polym = Number of polymorphic sites with a MAF equal to 1% or more. RMSE = Root Mean Square Error. The lower the RMSE the closer a given model is to the empirical observation. Here the 20 best models in terms of RMSE are displayed. Further details can be found in Supp. Table S3.

### Genetic differentiation and population admixture

Levels of genetic differentiation between each sampling location were computed using Weir and Cockerham’s Fst estimator θ (Weir & Cockerham, 1984) in vcftools v0.1.15 and the resulting values were used to construct a heatmap and a dendrogram in R using a custom script. Confidence intervals around Fst value were computed using Hierfstat with 1000 bootstraps. Isolation by distance (IBD) was tested between pairwise genetic distance as Fst/(1-Fst) (Rousset, 1997) and the riverscape distance using a Mantel test in R with 10 000 permutations.

Second, ancestry and admixture proportions were inferred for each individual using Admixture (Alexander et al., 2009) with K-values ranging from one to 25. Then the *snmf* function implemented in the LEA R package (Frichot & François, 2015) was used. Cross-validations were obtained using the cross-entropy criterion with 5% of masked genotypes. The default value for the regularization parameter was used to avoid forcing individuals into groups. Other model based clustering methods implemented in a DAPC (Jombart, Devillard, & Balloux, 2010) produced highly congruent results to those obtained using Admixture or LEA. Therefore, only the results obtained with Admixture are presented here. All admixture analyses were performed by keeping a single SNP per locus. Keeping either a random SNP or the SNP with the highest minor allele frequency produced similar results (not shown) and we only report the results of the analyses performed with SNPs showing the highest minor allelic frequency at each locus. We then correlated the individuals “stocking source” ancestry coefficient to individual levels of observed heterozygosity and verified the strength of the correlation using a t-test.

Finally, we used a modification of the fineSTRUCTURE package (Lawson, Hellenthal, Myers, & Falush, 2012) implemented in FineRADstructure (Malinsky, Trucchi, Lawson, & Falush, 2018) to infer levels of population genetic structure and ancestry from haplotype data derived from GBS data. Less than one percent of missing data were allowed. First, RADpainter was used to compute co-ancestry matrix, then individuals were assigned to populations with fineSTRUCTURE using 100 000 MCMC iteration for burn-in and the same number of sampled iteration with a thinning interval of 1000.

### Demographic simulations of historical stocking

Individual forward simulations were used to investigate the effect of recent stocking on admixture patterns and the genetic makeup of populations with a focus on the lower St. Lawrence River sampling sites only. The reason to focus on the St. Lawrence River is that data on stocking intensity was not always available for the tributaries. The goal of these simulations was to estimate a rate of effective migration from the sources of stocking to the St. Lawrence that would be necessary to explain the observed levels of admixture and hence help understanding the effect of stocking in this system. Here we focused on a restricted dataset in order to construct a simplified model of divergence made of three populations. The first population included sites from the different fluvial lakes and a river stretch separating lakes within the St. Lawrence River system, namely lakes Saint-Pierre (LSP), Saint-Françis (LSF) and Saint-Louis (LSL), as well as three different sampling localities within the river stretch between Montréal and Sorel and comprising Contrecoeur (MSC), Boucherville (MSG) and Varennes (MSV) sampling sites (*n* = 224 individuals, named SLR hereafter). The second population (lake des Deux-Montagnes, LDM *n* = 56) was separated from this first group based on patterns of admixture and shared ancestry (see results). Finally, the third population was the source of stocking populations and included as a single group the original site CHQ as well as JOS and TRE Lakes, (*n* = 100) which themselves originated from stocking with CHQ (confirmed by admixture and FineRADstructure analysis, see results). We grouped these individuals together for ease of simulations. Finally, the other source of stocking, PIG from the Kawartha system was not considered since no contribution to the admixture patterns in the St. Lawrence River was detected (see Results).

We simulated a neutral 10 000 kb long chromosome assuming a uniform mutation rate *μ* of 1e-8 per bp per generation and a recombination rate *r* of 1e-8 per bp per generation using slim v2.6 (Haller & Messer, 2017). First, we simulated an ancestral ideal population of size N = 2,400 for 80,000 generations to reach equilibrium. Then, the ancestral population was split into three different populations corresponding to the stocking source (SRC), the LDM and SLR with a reduced initial size of N = 800 (a range of different sizes were tested, see below). Given our observed data, we allowed for constant rate of migration between LDM and SLR with *m_SLR ->LDM_* = *0.0005* and *m_LDM →SLR ^=^_* 0.00025 (we explored a variety of parameters producing data closely similar to the empirical ones). We implemented asymmetric dispersal to reflect the expected downstream biased dispersal (e.g. Paz-Vinas et al. 2015; Rougemont et al., submitted) which was also observed in our data from LDM into SLR (see admixture results). Initially, no migration was allowed between the stocking sources (considered as a single unit based on patterns of shared ancestry) and either LDM or SLR. These populations kept diverging for 2,685 generations roughly corresponding to the postglacial divergence period (assuming a generation time of 6 years; Farrell et al., 2007), following the colonization of the study system. To reproduce the stocking event that was initiated about 15 generations ago, we introduced migration between the stocking sources and LDM, and SLR. Since more admixture was observed in LDM compared to SLR (see Results), migration into LDM was set 1.5 times higher than in SLR. We tested a range of migration rate from 1e-6 to 0.1. Then, after 82 685 generations, a set of individuals matching our empirical sample sizes were randomly sampled and exported into vcf format from which we computed a set of summary statistics corresponding to admixture levels, Weir & Cockerham F_ST_ and the numbers of polymorphic site above the 1% MAF. We also allowed for lower fitness of the stocking source population by implementing a mortality based filter (fitness callback in Slim v2, Haller & Messer 2017). We explored a set of fitness filters where 50, 100, 200, and 400 randomly chosen stocked individuals died, implying a mortality rate of 6%, 12.5%, 25% and 50% for N = 800. They allowed testing the idea that a small number of individuals effectively reproduced, but also considering various kinds of mortality-based filters such as lower local adaptation of those migrant individuals. Finally, we tested the effect of varying demographic scenarios through population increase or decrease by multiplying the size of each descendant populations by 2 (N = 1,600), 0.75 (N=600, closer to the estimated effective population size) and 0.5 (N=400). The ancestral population size was multiplied accordingly and the number of individuals that died was kept constant, implying a varying mortality rate. Simulations were repeated 50 times to assess the variability of admixture inferences. We then computed the root mean squared error (RMSE) for each scenario and computed the distance between the distributions of summary statistics and the empirically observed one, allowing us to compute how the model fitted the data. All scripts used with Slim are freely available and results can be reproduced using: https://github.com/QuentinRougemont/fwd_sims.

## Results

### Genetic diversity

An average of 3.23 million reads per individual was sequenced. Genetic diversity indices measured using haplotype data revealed a median Π value of 0.003. Median observed heterozygosity and gene diversity for polymorphic SNPs markers only were 0.144 and 0.148 respectively (Table 1). Two sites stood out as displaying significantly lower observed heterozygosity compared to all other sites, namely the populations from TRA (Ho = 0.088, Hs = 0.086, *Wilcoxon*-*test*, W = 759 *P* < 0.001) and PIG (Ho = 0.070, Hs = 0.066, *Wilcoxon*-*test*, W = 330, *P* < 0.001). Lower diversity for these two populations was also reflected by their Π estimates of 0.0016 and 0.0019 respectively (Table 1). Lakes of the St. Lawrence system (des Deux-Montagnes (LDM), Saint–Louis (LSL), Saint-Françis (LSF), and Saint-Pierre (LSP) Lakes) also displayed slightly lower diversity than the median diversity values (averaged Ho = 0.103, Hs = 0.105) whereas the opposite was observed for sites located in the fluvial stretch of the St. Lawrence between LSL and LSP (MSG, MSC and MSV; averaged Ho = 0.156, Hs = 0.161). The median proportion of polymorphic SNPs was 18% (s.d = 6%), with the least polymorphic site being the Ottawa River (OTT; 11%) which was expected given the small sample size of this sample (n = 7), whereas LSL was the most polymorphic location (31%). Ne estimates returned low values ranging from 14 (CHP) to 2308 (LSP) with four cases where the Ne value could not be estimated, either because of the paucity of polymorphic markers and/or limited sample size such as for OTT (result not shown). Pooling all Muskellunge from different sampling sites according to their inferred genetic cluster (see below) resolved this problem and revealed for instance a Ne value of 669 (95% CI = 493-823) for the St. Lawrence group (Table 1). It is worth noting that without correction for physical linkage all Ne values were downwardly biased, corroborating findings by Waples et al. (2016).

### Population differentiation

Global *F_ST_* estimate averaged over-all pairwise comparisons, was 0.211 (ranging from 0.0006 between LSL and MSC to 0.709 between TRA and PIG). Five geographically close locations within the St. Lawrence River (i.e. LSL *vs* MSG, LSL *vs* MSC, MSG *vs* MSC and both MSC and MSG *vs* MSV) displayed non-significant pairwise *F_ST_* values with confidence intervals overlapping with zero (Fig 2a; Table S1). When excluding all tributaries and inland lakes, *F*_ST_ values averaged over all sites from the Thousand Islands National Park (TIN) in the Upper St. Lawrence River downstream to LSP was 0.014. This value was driven by the higher differentiation of LDM when compared to all other sites, with an average *F*_ST_ of 0.036 (Fig2a). Indeed, the LDM site clustered separately from all sites located on the Upper and Lower St. Lawrence (Fig 2b). When removing the LDM site, the *F*_ST_ among all other sampling locations within the Upper and Lower St. Lawrence River dropped to 0.008, suggesting weak genetic differentiation and pronounced connectivity throughout the main stem of the St. Lawrence River. LSL, LSP, LSF and LSW displayed significant but weak *F*_ST_ (i.e < 0.009) between each other and between the three sites from the St. Lawrence section including MSC, MSG, and MSV. Accordingly, these sites clustered together in Fig 2b. The uppermost St. Lawrence population (TIN) displayed higher differentiation when compared to all other sites with *F*_ST_ ranging from 0.015 to 0.055. All tributaries were significantly differentiated from the St. Lawrence R. sites between LSL and LSP and displayed a much stronger level of differentiation than observed within the St. Lawrence R. itself with an averaged *F*_ST_ value between each tributary and all sites from the St. Lawrence R. being 0.20 for the Saint-Maurice R. (MAU), 0.119 for the OTT, 0.177 for the Chaudière R. (CHD), and 0.196 for the Yamaska R. (YAM). In the population tree topology, the CHD and MAU populations, both with presumably historically low number of muskellunge that were subsequently stocked, clustered close to the stocking source, yet their distinctiveness was strongly supported (Fig2b). The situation of Maskinongé L. (MSK), Ottawa R. (OTT) and Champlain Lake (CHP) was less clear as they appeared close to the source of stocking albeit the clustering was weakly supported in the tree (Fig 2b). There was a modest signal of IBD when all sites were included (mantel test r = 0.369 p = 0.0129) and considering only sites from the St. Lawrence R. (from TIN to LSP) revealed a much stronger and significant IBD pattern (r = 0.799, p = 0.0046, Supp Fig 2).

**Figure 2.**
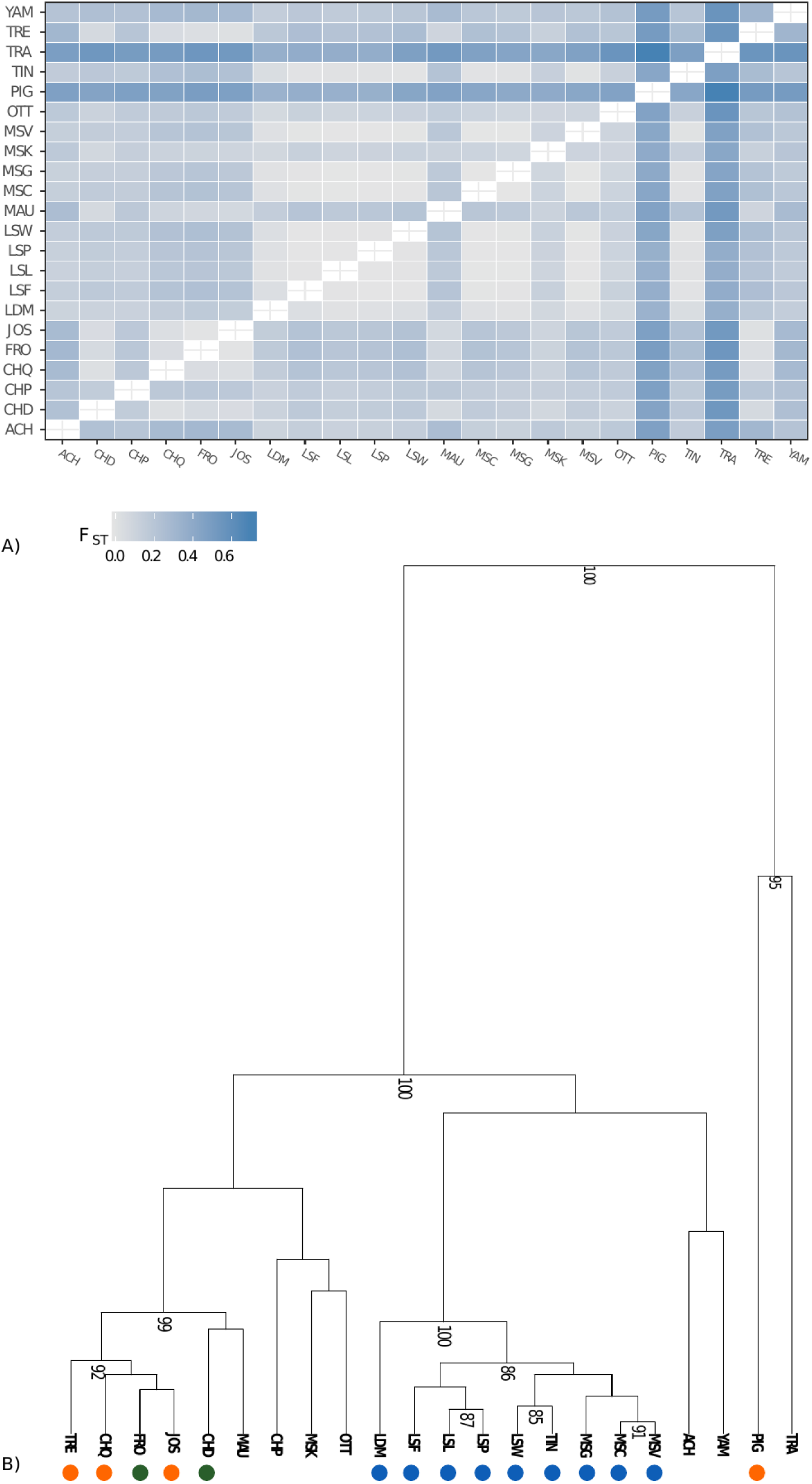
A) Heatmap of pairwise Fst values between each sample site. B) Clustering based on pairwise Fst values performed on each sample site. Only nodes with a bootstrap support higher than 80 are displayed. Orange dots = source of individuals used for stocking. Green dots = lake or river where Muskellunge were absent and have been introduced. Blue dots = site from the St. Lawrence River and des Deux-Montagnes Lake.

### Population Structure and Admixture

The analyses of admixture cross-validation consistently produced low cross-validation scores for K = 8 and K = 13 indicating that the number of clusters likely lies in between these values (Supp Fig S3). Similar results were obtained from LEA cross-entropy criterion with minimum values obtained around K = 12 and 13 while the DAPC provided the same results. For the sake of clarity, only results from the admixture analysis are detailed and presented in Fig 3a for K = 8 and 13 and Supp Fig S3 for K = 11 and 12.

**Figure 3.**
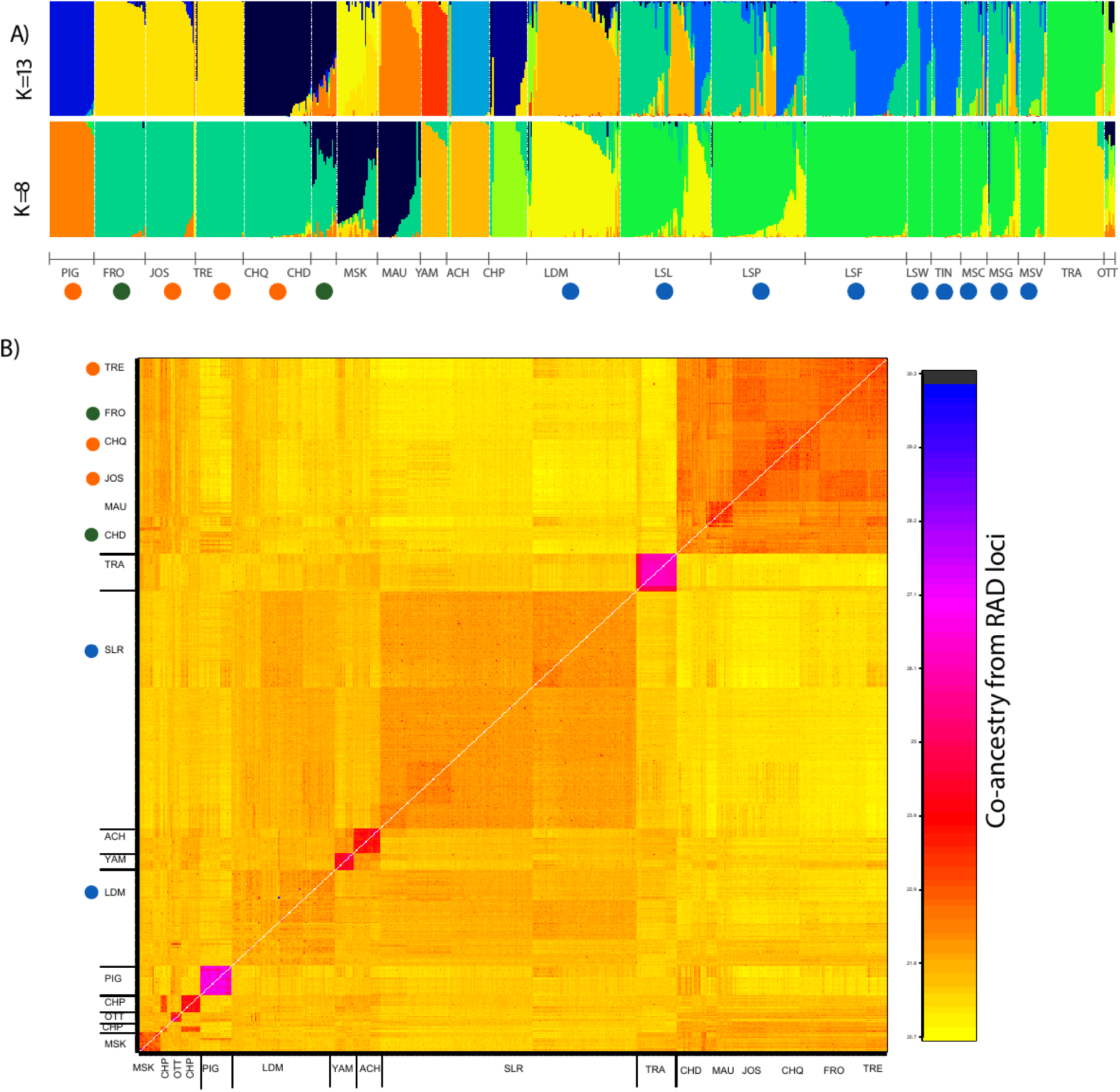
A) Levels of admixture for K = 8 and K = 13. Each color represents a distinct cluster and each bar represents an individual. See Fig1 for the labels of each dot in the graph; B) Co-ancestry matrix inferred by FineRADstructure. Each pixel represents individual co-ancestry coefficient inferred based on short haplotype loci. The labels summarize the major groups according to names of sampling sites. The higher values of coancestry coefficient sharing are depicted in darker colors whereas lower values of co-ancestry coefficient sharing appear in yellow colors.

Considering K = 8 revealed a clustering of the major groups used for stocking (i.e. CHQ, JOS, TRE) as well as the FRO introduced from the JOS population (mean q-value for the four lakes = 0.975; Fig 3). Here we considered individuals with 0.10 < q-value < 0.90 as being admixed, as commonly done in the literature (e.g. Valiquette et al. 2014, Létourneau et al 2018). Based on this criterion, we observed a complex pattern of population admixture in all tributaries and lakes outside the main stem of the St. Lawrence River. Thus, 13 fish (50%) from MAU displayed a membership probability > 0.10 of belonging to the source of stocking. More specifically, one individual was assigned to the “stocking” group with a q-value > 0.95 and the remaining individuals displayed mixed membership probability suggesting either backcrosses or advanced generations hybrids (q-values between 0.116 and 0.626). The 13 remaining individuals (50%) from MAU displayed a q-value > 0.90 of belonging to another group made of individuals from the MAU, CHD and MSK. All 15 individuals from CHD displayed mixed ancestry with membership probability of belonging to the “stocking” group ranging from 0.407 to 0.724. All Muskellunge from MSK were admixed (q-values ranging from 0.22 to 0.88). Among those, nine individuals (36%) displayed q-value of belonging to the “stocking group” ranging from 0.10 to 0.617 with no individual of apparent pure “stocking” origin (Fig 3a). Six (37.5%) and four individuals (57%) respectively from YAM and OTT were admixed (q-value > 0.10) with the stocking source (Fig 3a). None of these individuals was of pure stocking origin as they displayed q-values ranging from 0.10 to 0.211 and from 0.10 to 0.365 for YAM and OTT, respectively. In OTT, all seven sampled Muskellunge were admixed and none was assigned to a particular cluster. The Achigan R. (ACH) and YAM populations clustered together (mean q-value = 0.915) whereas TRA and CHP populations formed two separate clusters (q-value = 0.96 and 0.81 respectively). Finally, the Champlain L. showed signs of significant admixture from the stocking source for six individuals (30%) with 2 F_0_ immigrants and 5 introgressed individuals, the q-values ranging from 0.10 to 0.525.

Sites from the St. Lawrence River from the uppermost (TIN) to the most downstream site (LSP) tended to form a single group with the exception of LDM that clearly clustered separately (q-value = 0.84). In contrast with inland lakes and tributaries, we found weak evidence for admixture with the stocking source population. Thus, 21 individuals (8%) from the St. Lawrence R. (from TIN to LSP) displayed introgression with the stocking source (q-value ranging from 0.10 to 0.54) and no individual was of pure stocking origin. However, we observed more fish (n= 19, 33%) from LDM that were admixed with the stocking source (q-value ranging from 0.10 to 0.50) but none were of pure stocking origin. While there was limited admixture with the stocking source, we found evidence for admixture among fish from different locations within the system. Here, 6% of individuals from LSL were classified as F0 immigrants (q-value>0.9) from the nearby LDM population and 42% were admixed with LDM (Fig 3a). It is noteworthy that considering the spatial location of individuals in LSL (all collected fish within the SLR were georeferenced by GPS), we observed a tendency for the most admixed individuals (78%) to be preferentially found in the northern part of LSL whereas 87% of pure individuals were located preferentially in the southern part of LSL. Finally, 17 Muskellunge (30%) from LSP displayed admixture with LDM whereas 11 individuals (22%) from the St. Lawrence R. section including MSG, MSC and MSV were admixed with LDM (q-value > 0.10).

Considering K = 13 revealed the same general pattern of differential admixture whereby weak admixture with the stocking source was observed at all sites within the main stem of the St. Lawrence R., a somewhat more pronounced admixture in LDM and the highest admixture being observed in tributaries and inland lakes (except FRO where no Muskellunge occurred before stocking) (Fig. 3a). However, K=13 revealed more separation between some population clusters. Thus, Muskellunge from YAM and ACH tributaries were clearly assigned to two different clusters corresponding to their local river. The original stocking source (CHQ) clustered independently (mean q-value = 0.95) of the derived individuals of JOS-TRE stocking sources that grouped together (q-value =0.96). While the FRO lake still clustered with JOS, fish from MAU clustered independently (q-value = 0.87) but with one individual showing a F_0_ immigrant profile (q-value > 0.95) from the JOS-TRE stocking group and seven other individuals showing signs of introgression (0.10 < q-value < 0.53) (Fig 4a, b). Five Muskellunge (20%) from MSK displayed a q-value > 0.90, while 95% of the remaining individuals showed introgression (0.13< q-value < 0.730) with the JOS-TRE stocking source (Fig 3a, Fig4). The CHP site tended to form a separate cluster (mean q-value = 0.81) but with one F_0_ immigrant from CHQ stocking source and five individuals (21%) showing admixture (F1 like profile) from the JOS-TRE group (Fig 3a). Individuals from the St. Lawrence River from TIN to LSP were now separated into two admixed groups. In particular, individuals from the uppermost site TIN were assigned to this cluster (mean-q value = 0.87) while 31%, 26%, 12% and 7% of individual respectively from LSF, LSW, LSL and LSP were now assigned (q > 0.90) to this new group. In these same sites, respectively 31%, 27%, 22% and 37% of individuals, as well as 3.5% in LDM were admixed (Fig 3a, top panel). Finally, five individuals (9%) from LSL were assigned as putative F0 immigrants from LDM and 16 individuals (29%) displayed mixed membership probability. The same was observed in LSP with no F_0_ immigrants but 15 individuals (26%) admixed with LDM. Overall, Muskellunge from the St. Lawrence displayed lower stocking q-value membership than the LDM or tributaries and inlands lakes (SLR range: 0.10 to 0.63, LDM range: 0.10 to 0.51, tributaries range: 0.10 to 0.99, see details and boxplots in Fig 4a). Similarly, the number of admixed individuals was lower in the St. Lawrence (8%) than in the LDM (31%) and the different tributaries and inland lakes (42%) as detailed in Fig 4b. Finally, there was a global significant and positive correlation between the level of stocking ancestry and the observed heterozygosity for K = 13 (rho spearman = 0.37, t = −15.578, df = 452.03, p-value < 2.2e-16) as well as for K = 8 (rho spearman = 0.26, t = 12.863, df = 513.38, p-value < 2.2e-16).

**Figure 4.**
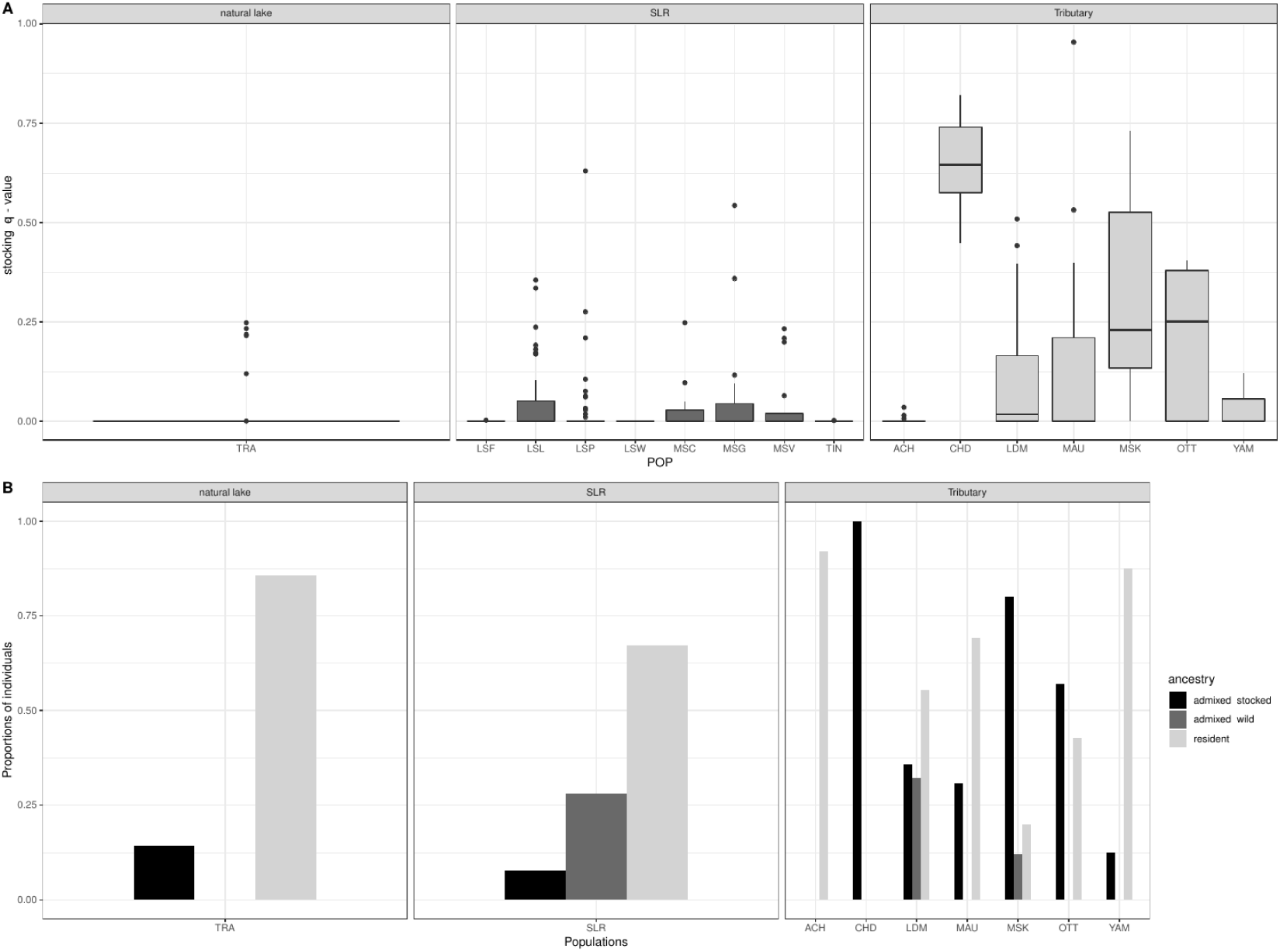
A) Boxplot of “stocking source” ancestry coefficient (q-values) in the single non supplemented population and in the different supplemented populations separated according to whether they occur in the St. Lawrence River itself (SLR) or its tributaries. B) Proportion of individuals assigned to different classes of ancestry: individual assigned as “resident” if q-value > 0.9, individual assigned as of “admixed domestic ancestry” if q-value of belonging to a “stocking source cluster was above 0.1, individual assigned as of “wild ancestry” if q-value of belonging to another foreign population above 0.10.

Results from co-ancestry analyses at the haplotype level provided additional information about ancestry between individuals and revealed patterns consistent with admixture analyses at different K values highlighting the similarity observed at K = 8 and the finer differences for K = 13. In particular, two major blocks stood out corresponding to *i)* the populations used for stocking and *ii)* the St. Lawrence R. Here, the CHQ, JOS and TRE sources as well as the derived populations FRO and CHD appeared closely related (top panel in Fig 3b). All individuals from MAU, 24% from MSK and 9% from CHP shared ancestry with these source populations (Fig 3b). The second major block was made of individuals from the different sampling sites from the St. Lawrence River (TIN to LSP excluding tributaries) and displayed moderately shared co-ancestry with LDM. In particular, 36% of LSL, 12% of LSP and 16% of the MSC, MSG and MSV group of individuals displayed closer co-ancestry to LDM than to remaining individuals from SLR. One individual from JOS displayed close ancestry to the St. Lawrence River fish as observed in admixture analyses. The seven individuals from OTT were closely related to a group of five individuals (9%) from LDM, indicating possible downstream dispersal from the Ottawa River to LDM or a common ancestral origin. All fish from ACH and YAM were well separated (K = 13 in admixture), but displayed close ancestry, highlighting their close relationship (K = 8 in admixture). Finally, both PIG and TRA individuals formed well-separated clusters of ancestry (Fig 3b).

Altogether, our analysis of genetic differentiation, admixture and fineRADstructure suggest the existence of at least eight major groups, the first group corresponding to the “source, introduced or stocked populations” made of CHQ, JOS, TRE, FRO, CHD, MSK L. and MAU R., a second cluster composed of the ACH R., a third cluster corresponding to the YAM R., a fourth group comprising the whole St. Lawrence R. (TIN to LSP), and different independent groups corresponding to LDM, TRA, and PIG and the last group being the CHP L. We avoid classifying the OTT R. in any group due to caveats associated to its small sample size. Within the St. Lawrence R., it is possible to further differentiate (but with high caution as discussed below) an “upper group” composed of TIN, LSW, and LSF and a lower group composed of all sites of the Lower St. Lawrence downstream of LSF. Finally, depending on the clustering considered, MSK and the MAU could also be further separated into two different clusters, yet introgressed by fish from stocking sources.

### Simulations of admixture

A total of 17 different migration rates were simulated with population sizes of Ne = 400, 600, 800 and 1,600 to generate expected admixture values following the scenario detailed in Supp Fig S4 and Supp Table S2. Results indicated that over all scenarios, the one with N = 600 (closer to estimated Ne in the SLR) systematically displayed lower RMSE (root mean squared errors) than those with higher Ne (Supp Table S4). For instance, only N = 600 resulted in a similar level of polymorphism (i.e. polymorphic SNPs at the 1% level) as our empirical data, while all other datasets were significantly different in terms of number of polymorphic sites. Therefore, for the sake of simplicity, we only presented results for the N = 600 scenario (the 20 best model related to N =600 are presented in Table 2); other scenarios for models with the lowest RMSE are presented in Supp. Table S3). The best scenarios in terms of RMSE were those in which migration rate from the stocking source (SRC) to the St. Lawrence (LSW to LSP) ranged between *m* = *0*.*0025* and *m* = *0*.*005*. Migration rate was systematically higher from the SRC to LDM with values ranging from *m* = *0*.*00375* to *m* = *0*.*0075* (ms M20 to M23, similar to M9 to M11 but includes mortality filters; Table 2). None of the scenarios in which the migration rate was set higher (from *m* = *0*.*01* to *m* = *0*.*30*) provided a good fit (Table 2, Supp Table S3). In particular, M16 theoretically would reproduce most closely the expected level of migration (m ~ 0.5) due to the intensity of stocking, but poorly fitted the data (Suppl. Table S3). Scenarios M1 to M8, with lower migration rates (*m* < *0*.*001*), did not properly reproduce observed data.

While the level of genetic differentiation between SRC and SLR or between SRC and LDM were qualitatively close to those observed empirically in models M20 to M23 (simulated *F*_ST_ range ~ 0.192 to 0.298, Table 2), none of the scenarios were able to reproduce the divergence between SLR and LDM (mean *F*_ST_ for the 20 best scenarios = 0.011 versus 0.036 in our data set; Table 2). The difference between simulated and observed differentiation was significant (p < 0.005), meaning that some non-modeled processes such as a long isolation period or a smaller *Ne* in LDM or that two populations originating from evolutionary divergent lineages may have contributed to the divergence between the two populations. Overall, scenarios with the mortality-based filter produced the lowest RMSE, with the best scenario (RMSE = 0.05) being one where 50 individuals randomly died (Table 2). However, scenario M9 where no individual died also produced a low RMSE of 1.60. Several scenarios with mortality of 100 to 400 individuals presented intermediate RMSE values (Table 2), indicating that varying rates of mortality for N = 600 could explain the data. Increasing the mortality rate to 500 individuals (i.e. 83%) resulted in RMSE of 1.73 and fitted well the data (not shown).

## Discussion

The primary objective of this work was to assess the effect of extensive fish supplementation from genetically differentiated groups of Muskellunge on the genetic diversity and population structure in the St. Lawrence River, the majority of its main tributaries as well as in Québec inland lakes, Canada, where the species is found. To do so, we first characterized Muskellunge genetic structure and diversity in a wide variety of sites, including the main sources of admixture. Second, to understand the contribution of these supplementation efforts in generating admixture patterns in the St. Lawrence R. system, we compared our data to the predictions obtained from simplified demographic scenarios.

### Genetic structure between distant lakes and isolation by distance within the St. Lawrence River

A common issue in population genetics is the delineation of population structure that can be confounded with the isolation by distance pattern, in which genetic differentiation increases with geographic distance due to the action of genetic drift when dispersal is limited (Wright, 1943). Indeed, patterns of isolation by distance are pervasive across various species (Meirmans, 2012; Sexton et al., 2014). This can result in false inference of population structure as commonly used algorithms partition continuously distributed variations among samples into discrete clusters (Frantz et al. 2009; Meirmans, 2012). Here, our data indicated the occurrence of both clusters of genetically highly differentiated groups (i.e. among distant and isolated water bodies) and of significant isolation by distance within the St. Lawrence River accompanied by relatively weak population structure.

Regarding the major clusters, the combination of *F*_ST_ values, admixture and fineRADstructure analysis indicated that the most divergent and least polymorphic populations were the most geographically isolated ones (TRA and PIG). The two major population groups were *i)* the St. Lawrence River comprising all sites from TIN to LSP with clear patterns of shared co-ancestry and *ii)* the source of stocking represented by CHQ, its derived populations (i.e. JOS, TRE), and the sites where Muskellunge was introduced using those source populations. The MAU River, which has no recorded history of stocking to our knowledge, clustered with the stocking source suggesting the possible occurrence of migrants from further upstream stocked waters in this watershed. All analyses indicated a complex admixture pattern in MSK and OTT with for instance 32% of individuals from MSK displaying close ancestry to the stocking source. While this suggests a relatively pronounced genetic impact of stocking, interpreting those results must be done cautiously as small sample size can cause biases that translate as a “mix of multiple drifted groups” (Lawson, et al., 2018). The findings of only a modest IBD in the complete dataset and rather strong signal of population differentiation and structure outside the St. Lawrence R. suggest that populations from different inland lakes and tributaries, isolated by the presence of dams and natural barriers, are evolving mainly through drift, echoing previous findings on Muskellunge in other geographic areas (Turnquist et al., 2017). Over the whole data, the strength of IBD is likely diminished by a combination of long distance natural dispersal, the non-linear distance among isolated lakes and rivers, as well as stocking events over a large distance (Miller et al., 2017).

On the other hand, the clear distinction from the stocking source populations, the linear relationship between distance and the significant population differentiation within the St. Lawrence River support the hypothesis that the original genetic structure has weakly been affected by stocking as in other parts of the species range (Miller et al. 2012). Similar findings of IBD were observed in the Great Lakes based on microsatellite data (Kapuscinski et al., 2013; Miller et al., 2017). The behavior of the muskellunge, especially spawning site fidelity and possible natal homing (Crossman, 1990; Jennings, Hatzenbeler, & Kampa, 2011; Margenau, 1994; Miller, Kallemeyn, & Senanan, 2001) are two factors that can contribute to both IBD and the establishment of population structure. While individuals from the St. Lawrence River display shared co-ancestry from the uppermost to the lowermost sites, admixture for K = 13 however, splits the cluster in two different groups with over 72% of individuals from TIN being assigned to this new cluster and a decreasing number of individuals being assigned to this cluster when moving downstream. Such patterns suggest higher genetic drift in the upstream site, and most likely reflects the idea that restricted dispersal generates increasing differentiation with increasing distance, resulting in an overestimation of the number of clusters by the clustering algorithm (Meirmans, 2012, Bradburd, Ralph & Coop, 2018). While this artefact represents mostly the decay of ancestry with increasing distance from our most upstream site, it is likely that the two major dams present on the St. Lawrence (Fig1) now constrain upstream gene flow (towards LSF, LSW and TIN). For instance, no gene flow from LDM to LSW was observed whereas downstream biased dispersal is expected (see below). Such downstream biased gene flow is further expected because of the unidirectional movement of water and its effect has been demonstrated across various fish species (Paz-Vinas, Loot, Stevens, & Blanchet, 2015, Paz-Vinas et al. 2013, Morrisey et de Kerchove, 2013, Rougemont et al., Submitted).

We observed a variable proportion of fish from sites between LSL and LSP displaying a non-negligible amount of ancestry profile resembling those of the LDM (itself genetically divergent from the St. Lawrence River) suggesting downstream biased dispersal. No migrant from LDM was inferred in the upstream LSF as expected due to the impassable Beauharnois-Les Cèdres hydropower dams (Fig1). The dispersal from genetically divergent background resulted in admixture of LSL with LDM and a signal of higher admixture with LDM on the northern shore of LSL. Interestingly, a contrasted population genetic structure between the northern and southern shores of LSL was also previously reported in the Northern Pike (Ouellet-Cauchon, Mingelbier, Lecomte, & Bernatchez, 2014) and Yellow Perch *Perca flavescens* (Leclerc, Mailhot, Mingelbier, & Bernatchez, 2008) whereby in both species, fish from the south shore of the LSL were more similar to those elsewhere within the St. Lawrence R. than those from the north shore. It is noteworthy that this lake is characterized by contrasted water masses stemming from the St. Lawrence R. on the south shore (so-called green waters) and water from the Ottawa River on the north shore (so-called brown waters) (Hudon et Carignan, 2008; Leclerc, Mailhot, Mingelbier, & Bernatchez, 2008). Interestingly, our forward simulation did not allow to reproduce the pattern of genetic differentiation observed between LDM and the group of populations from the SLR, indicating that the modelled intensity of effective migration between the two groups were still too high to properly reproduce the observed weak divergence. Given the above, it is likely that various selective processes are acting to reduce effective migration in the system. Therefore, it would be relevant to distinguish the respective contribution of ecological (e.g. temperature, pH, light extinction, etc.) and historical (e.g. glacial refugia) factors responsible for this repeated pattern of divergence observed between the north and south shores of LSL among different species. One could also take advantage of our forward simulation framework to construct a more complex model that includes selection against migrant, secondary contacts and biased dispersal. This would allow moving beyond the simple description of patterns, but was beyond the goal of our study.

### Stocking preferentially impact smaller populations

Correlation between stocking intensity and admixture have been previously documented in a range of species, but mainly in salmonids (Campos, Posada, & Morán, 2008; Finnegan & Stevens, 2008; García-Marín, Sanz, & Pla, 2006; Perrier, Baglinière, et al., 2013; Perrier, Guyomard, et al., 2013; Sønstebø, Borgstrøm, & Heun, 2008) with an emerging general pattern reflecting a tendency toward decreasing admixture proportion after stocking cessation (Hansen et al., 2009; Harbicht, Wilson, & Fraser, 2014; Létourneau et al., 2018.; Perrier, Baglinière, et al., 2013; Valiquette, Perrier, Thibault, & Bernatchez, 2014). Our data suggest a complex relationship regarding admixture proportions and stocking. Indeed, the most extensively stocked populations (Lower SLR) displayed the lowest amount of stocking ancestry suggesting that since the cessation of stocking in 1998, the majority of chromosomal blocks introgressed from the stocking source populations have been removed from the genome, just as in other geographic areas (Miller et al. 2012) and were not detectable by our approach. Such patterns are expected if stocked individuals display low survival, low reproductive success (hence resulting in little introgression), and/or if selection is acting efficiently against introgression, as supported by several of our models in which 400 individuals would die. In contrast, other supplemented tributaries and lakes (e.g. MAU, MSK, LDM, OTT) clearly displayed signs of shared ancestry with the stocking source. A likely hypothesis is that the higher effective population size and large-scale connectivity existing in the Lower SLR played a critical role in the efficacy of selection against introgression (Glémin, 2003). On the other hand, genetic drift might overcome selection against introgression in smaller, isolated populations (Frankham et al. 2010) leading to fixation of “foreign” alleles and higher footprints of admixture. Such variable outcomes of stocking on admixture have been documented in salmonids (Perrier, Guyomard, et al., 2013). More specifically, Perrier et al. (2013) found high rates of admixture in weakly stocked populations as well as a heavily stocked population in which wild individuals were still present.

Here, using forward simulations, we investigated which levels of migration from the stocking source to the SLR and LDM would generate similar patterns of admixture and genetic differentiation. Our simulations indicated low levels of migration (*m* < 0.005) and high mortality that encompasses both low survival and/or reproductive success of stocked fish, as observed in other species (Araki et al., 2007, 2009; Araki & Schmid, 2010; Christie, Ford, & Blouin, 2014; Thériault, Moyer, & Banks, 2010 Thériault, Moyer, Jackson, Blouin, & Banks, 2011). This result indicates that the supplementation effort does not translate into effective migration in the long term, which further supports our hypothesis that introduced individuals may have lower fitness than fish from the resident local populations. Lower fitness was expected given that maladaptive change that reduces fitness in the wild can be observed within just one captive generation (Araki et al, 2007, 2009, Christie et al. 2012; Fraser et al. 2018). Here, eggs from adults were kept and reared in a hatchery fish farm and more than half of them were released as juveniles (5-27 cm in length, De LaFontaine, unpublished) so that selected traits in the hatchery can differ from those providing highest fitness in the wild (Frankham, 2008). Moreover, this effect may have been amplified by the use of genetically divergent fish. Therefore, thanks to appropriate modeling, our results give support to the idea that stocked fish from ecologically and genetically divergent populations display lower success than their wild relatives in the St. Lawrence. Similarly, Perrier et al. (2013) found a 10-25 times lower survival of stocked fish relative to wild fish. These results are not entirely comparable since they used a higher migration rate (~0.06 to 0.24), whereas, in our case, a low migration rate was found to better reflect the inefficiency of stocking. Despite its ease of implementation, to the best of our knowledge, forward simulations have rarely been used to investigate the effect of stocking (but see Perrier et al., 2013, Fernandez-Cebrian et al., 2014). Further, coalescent simulation (e.g. Hudson, 2002) could be used to quickly generate variation data and couple with our forward simulations. In this way, both neutral processes and realistic demographic hypotheses incorporating positive and negative selections could be coupled together. These data could be analyzed in an Approximate Bayesian Computation framework (Beaumont et al., 2002) to test more realistic scenario of migration and subsequent admixture reflecting complex process in nature. Therefore, while our model provided a simple means of assessing the rate of effective migration, further effort would be critical toward obtaining more biologically meaningful simulation reflecting complex process in nature over spatial and temporal scales. Finally, it should be emphasized that a potential limitation of admixture software is that it may fail to detect admixture in the St. Lawrence River if admixture events and drift affect all samples in the same way (Lawson et al., 2018). Therefore, even accurate forward modelling could be affected by inaccurate estimates of co-ancestry that might miss localized genome-wide introgression. We further stress that admixture analysis, as a global ancestry inference method may have failed to detect small tracts of introgression that potentially have not been purged (e.g. Leitwein et al. 2018). A local ancestry approach (e.g. Dias-Alves et al. 2018) would be required but given the relatively small density of SNPs available here this was not possible. The putative positive and negative fitness consequences of any remaining alleles of stocked origin would therefore require further investigations.

### Does stocking reduce genetic diversity in large populations?

Levels of population genetic diversity were generally modest, which is consistent with previous studies on Muskellunge based on microsatellite data in other geographic areas (Miller et al., 2012; Wilson et al., 2016, Turnquist et al., 2017, Miller et al., 2017). Although the various biases associated to RADseq like data may complicate the interpretation of genetic diversity estimates (Arnold, Corbett-Detig, Hartl, & Bomblies, 2013; Cariou, Duret, & Charlat, 2016; Gautier et al., 2013), some general patterns emerged from our data. First, the population with the lowest level of genetic diversity was also the one that has historically never been stocked (i.e. TRA), a difference that can best be explained by the small size and strong isolation of this lake restricting opportunity for gene flow and implying small effective population size. Giving the species life history traits, in particular a long life span (~ 30 years, Casselman, Robinson, & Crossman, 1999), as well as the territorial behavior of this predator (Becker, 1983), small effective population sizes in such isolated populations are expected (Romiguier et al., 2014), as previously reported for the Muskellunge’s sister species the Northern Pike (Miller & Kapuscinski, 1997). Second, there was a significant and positive correlation between stocking ancestry and genetic diversity. Stocked populations from the Lower SLR and LDM, which have been the most extensively stocked (> 630 000 fish released), displayed lower genetic diversity than the remaining supplemented tributaries and inland lakes, or than introduced populations from CHA, TRE, JOS, and FRO. Moreover, the most extensively stocked populations were those showing the lowest stocking ancestry. In Muskellunge, Scribner et al. (2015) found that stocked populations displayed higher genetic diversity when stocking was done using multiple strains. Similarly, Valiquette et al. (2014) and Ferchaud et al. (2018) found that genetic diversity of the stocked populations in Lake Trout (*Salvelinus namaycush*) was higher than unstocked populations. Given the fact that the most extensively stocked Muskellunge population (St. Lawrence River) is the one that showed the smallest level of domestic ancestry and lowest levels of genetic diversity amongst all stocked populations, we hypothesized that selection against introgression may have been more efficient to remove non-native alleles of stocked fish with a strongly divergent genetic background. Such hypothesis is supported by findings in other species. In particular, Hansen et al. (2009) found that Danish Brown Trout (*Salmo trutta*) that were stocked for 60 years were subject to selection against non-native alleles of stocking origin. Similarly, Muhlfeld et al. (2009) found a decline in fitness of Cutthroat Trout (*Oncorhynchus clarkii*) crossed with non-native Rainbow Trout (*O*. *mykiss*) over generations. Therefore, our results indicate that the intensity of stocking alone may be a poor predictor of the extant of admixture in a given population and further investigations of the fitness effect of admixture and introgression is needed. In particular, identifying deleterious mutations would be necessary to quantify the mutation load (e.g. Ferchaud et al. 2018) and, combined with our simulation framework could help to refine our understanding of the fitness effect of stocking on wild populations.

### Conservation applications

Knowledge of the genetic structure of fish populations is essential for proper and sustainable stock management since it allows identification of groups of individuals that are genetically and often geographically unconnected, therefore implying at least some demographic independence. Our results indicate that Muskellunge samples studied here can be separated into six major population groups. First, the Upper and Lower St. Lawrence River (Thousand Island National Parks to Saint Pierre L.) could be considered as a single group characterized by a pattern of isolation by distance in which ancestry decays with genetic distance between individuals. This large scale pattern should be considered in management decisions, especially considering that on the one hand, upstream movements between the Lower and Upper sections of the river are restricted by large dams but in the other hand, these dams do not apparently prevent gene flow from upstream to downstream. The second group is made of individuals from des Deux-Montagnes L., showing a clear shared co-ancestry with those of the Saint-Lawrence River, especially with Northern Saint-Louis L. and showing a downstream biased dispersal from des Deux-Montagnes L. to Saint-Louis L Saint-Pierre L., and to the Saint Lawrence R. stretch including MSG, MSV and MSC. The third group comprises tributaries draining into the St. Lawrence River. Each tributary appears as genetically distinct units with some nuance depending on whether the Muskellunge was naturally present or not. In particular, Yamaska R. and Achigan R. did not display any sign of stocking source introgression whereas Ottawa R. showed some extent of shared ancestry with the stocking source. On the other hand, historical data (Vézina, 1977) and our genetic results suggest that Muskellunge was absent or in very low abundance in Chaudière R., and Saint -Maurice R., explaining why they both showed a shared ancestry with the stocking source despite being geographically distant. The fourth group is represented by lakes where Muskellunge has been introduced (Joseph L., Tremblant L. and Frontière L.), exhibiting pronounced genetic similarity with the Chautauqua source of stocking. The fifth group includes stocked lakes where Muskellunge was initially present (Maskinongé and Champlain lakes) with each of these lakes representing a genetically distinct population. Finally, Traverse Lake represents the sixth group corresponding to a wild unstocked population with a unique genetic background. In isolated lacustrine and river systems originally unoccupied or with presumably low Muskellunge abundance, stocking efforts appear to have been successful in enhancing population and contribute to support angling activity. In the lower St. Lawrence, in St.Louis L., Dumont et al. (1991) also made a similar conclusion following an analysis of data from the first half period of the stocking program. These authors have shown that the two most important variables explaining the muskellunge year class strength in LSL were the number of stocked fish in the lake in a given year and the year class strength of the year before (Dumont et al. 1991). However, in contrast, admixture and simulations results suggest that extensive stocking in the lower St. Lawrence River has been inefficient and did not contribute significantly in sustaining local Muskellunge stocks in the long term. We therefore recommend avoiding this practice in the future without a good knowledge of the initial stock abundance, genetic diversity, structure and level of genetic connectivity prior to stocking as these are key parameter to ensure the success of any future stocking program. In cases where supplementation is necessary (e.g. in Thousand Islands massive die-off recently occurred due to viral hemorrhagic septicemia) we advocate the use of local brood sources and to avoid long term maintenance in captivity. Here it is worth noting that sampling was heterogeneous as composed of individuals both caught during spawning and by anglers during their feeding activity. Ideally future studies should aim at sampling mostly spawning individuals in order to be less biased by the presence of dispersers that may not be contributing to gene flow within a given sampling site.

In conclusion, we found that Muskellunge populations were spatially structured into a set of different groups, whose differentiation was affected by the levels of stocking, natural connectivity or geo - graphic isolation. Our results will be useful to identify the appropriate spatial scale at which fishery manage - ment and habitat protection measures should be applied. We also found that the genetic consequences of stocking on admixture and genetic diversity affected differentially Muskellunge from the St. Lawrence River, its tributaries and inland lakes. Disentangling the long-term evolutionary consequences of this history of stocking would require further investigations from denser genome wide data.

## Acknowledgements

We express our gratitude to Québec’s muskies anglers who collected most of the samples, especially Marc Thorpe, Mike Lazarus and Mike Phillips. We thank Christopher Legard (New York State Department of Environmental Conservation) for collecting and sharing Chautauqua Lake samples. We thank Samuel Cartier for collecting Lake Champlain fish. Thank to Chris Wilson (Aquatic Research and Monitoring Section Ontario Ministry of Natural Resources and Forestry) for sharing Pigeon Lake DNA samples and Christopher Wilson (Fish Culture SectionOntario Ministry of Natural Resources and Forestry) for sharing hatcheries and stocking history between Ontatio and Québec. Thank to Chris Wilson (Aquatic Research and Monitoring Section Ontario Ministry of Natural Resources and Forestry) for sharing Pigeon Lake DNA samples. We also thank Nicolas Auclair, Florent Archambault, Rémi Bacon, Christian Beaudoin, Anabel Carrier, Chantal Côté, Julie Deschesnes, François Girard, Guillaume Lemieux, Louise Nadon, Yves Paradis, Geneviève Richard, and Éliane Valiquette for logistic, field and laboratory assistance. Funding was provided by the Ministère des Forêts, de la Faune et des Parcs du Québec and the Canadian Research Chair in Genomics and Conservation of Aquatic Resources.

## Data availability

Raw sequencing data will be deposit on NCBI.

